# Human sleep spindles track experimentally excited brain circuits

**DOI:** 10.1101/2025.01.07.631687

**Authors:** Jude L. Thom, Bernhard P. Staresina

## Abstract

Spindles are hallmark oscillations during non-rapid-eye-movement (NREM) sleep. Together with slow oscillations (SOs), they are thought to play a mechanistic role in the consolidation of learned information. The quantity and spatial distribution of spindles has been linked to brain activity during learning before sleep and to memory performance after sleep. If spindles are drawn to cortical areas excited through pre-sleep learning tasks, this begs the question whether the spatial distribution of spindles is flexible, and whether their regional expression can also be manipulated with experimental brain stimulation. We used excitatory transcranial direct current stimulation (tDCS) to stimulate the left and right motor cortex in a repeated-measures experimental design. After stimulation, we recorded high-density electroencephalography (EEG) during sleep to test how local stimulation modulated the regional expression of sleep spindles. Indeed, we show that excitatory tDCS of local cortical sites before sleep biases the expression of spindles to the excited locations during subsequent sleep. No effects of localised tDCS excitation were seen for SOs. These results demonstrate that the spatial topography of sleep spindles is neither hard-wired nor random, with spindles being flexibly directed to exogenously excited cortical circuits.

**Graphical Abstract.**
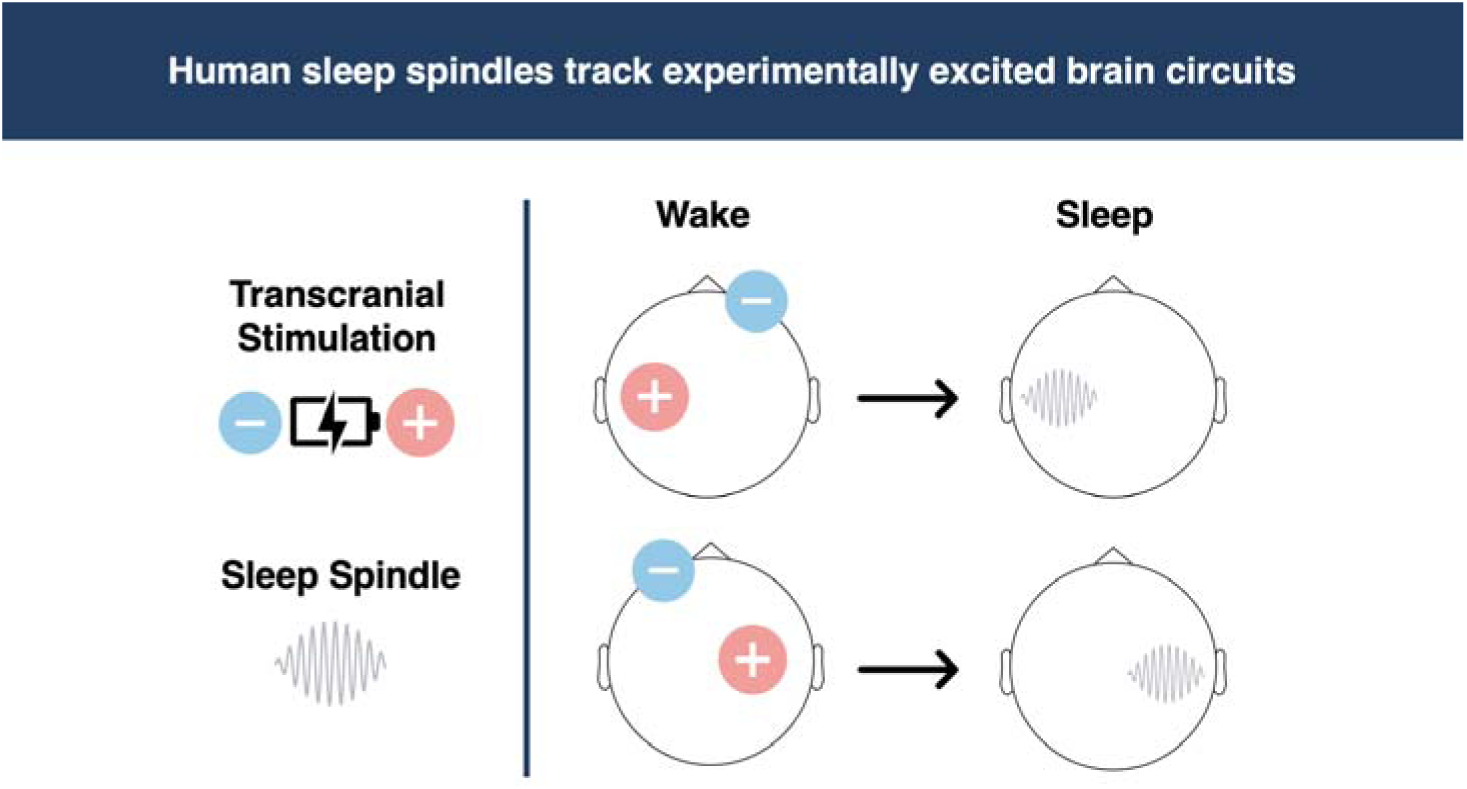

**Statement of Significance:** Spindles are signatures of NREM sleep linked to memory consolidation, intelligence, and neurological disorders. Recent work shows the spatial distribution of spindles mirrors cortical activation patterns during learning. If spindles are preferentially expressed at cortical sites engaged during prior waking, the question arises whether spindles can be directed by inducing excitation exogenously with non-invasive brain stimulation. Here, we applied anodal-tDCS to either left or right motor areas before a nap. Indeed, we found that spindle (and not SO) rates are greater at cortical sites excited by tDCS. This result suggests the spatial distribution of spindles is flexible and is sensitive to experimental excitatory stimulation. This has implications for targeted rehabilitation of specific cortical areas and for developing sleep-based brain-computer interfaces.

## Introduction

Sleep spindles are waxing and waning oscillations (12-16 Hz) generated in cortico-thalamic loops and expressed across the cortex during NREM sleep (Andrillon et al., 2011; Fernandez & Lüthi, 2020; Mak-McCully et al., 2017; Sarasso et al., 2014; Staresina, 2024; Staresina et al., 2015; Steriade, 2006). SOs are high-amplitude ∼1Hz oscillations which reflect increases and decreases in the excitation of larger neuronal populations (Massimini et al., 2004; Steriade, 2006; Vyazovskiy & Harris, 2013). Together, spindles and SOs are believed to play a key role in the synaptic plasticity and inter-regional communication necessary for memory consolidation (Diekelmann & Born, 2010; Ngo et al., 2020; Niethard et al., 2018; Schreiner et al., 2021; Staresina et al., 2015). Biophysical experiments indicate that spindles facilitate the influx of calcium into synaptic dendrites, which sets the process of synaptic potentiation in motion (Chittajallu et al., 1998). For instance, inducing artificial spindle-like activity with intracellular electrical stimulation increased calcium levels (Rosanova & Ulrich, 2005), while the power of spindles correlated with increased dendritic calcium activity (Seibt et al., 2017). Besides facilitating conditions for synaptic plasticity, spindles also provide time windows of heightened hippocampal-cortical communication (Ngo et al., 2020), with recent studies showing they enhance the precision of neural co-firing at frequencies optimal for of spike-timing-dependent-plasticity (Dickey et al., 2021; Staresina et al., 2023). These properties make spindles uniquely suited for transmitting information across neural populations and consolidating it through synaptic plasticity. Note that we refer to fast-spindles as “spindles” here, distinguished from slow spindles, which have a slower frequency profile (8-12 Hz) and have not been linked to memory processing (Mölle et al., 2011).

The ability to synchronise memory systems and induce plasticity suggests that spindles may play a mechanistic role in transferring information across memory systems and stabilising memory traces (Diekelmann & Born, 2010; Fernandez & Lüthi, 2020; Staresina, 2024; Ulrich, 2016). Indeed, the quantity, amplitude, and duration of spindles have been linked to behavioural improvements in memory after sleep (Bergmann et al., 2012; Cairney et al., 2018; Gais et al., 2002; Holz et al., 2012; Latchoumane et al., 2017; Mednick et al., 2013; Schabus et al., 2004; Schreiner et al., 2021). However, spindles are not uniformly distributed; instead, they display a spatial topography which varies with cognitive demands during prior wake (Clemens et al., 2005, 2006; Cox et al., 2014; Nir et al., 2011; Petzka et al., 2022; Piantoni et al., 2017; Tamaki et al., 2013). This topographical flexibility suggests that the local expression of spindles is impacted by pre-sleep task engagement. Recently, Petzka and colleagues investigated whether spindle topographies would adapt based on task demands (Petzka et al., 2022). They found that spindle expression mirrored the topography of engagement (expressed in alpha/beta power decreases) observed during a learning task before sleep. The overlap between spindle topographies and the distribution of task-induced alpha/beta power effects was predictive of memory retention after sleep. Of note, previous work has shown that decreases in alpha/beta power are related to increased cortical excitability, to increases in gamma power (>40 Hz), and to increases in neuronal firing-rates (Bonnefond & Jensen, 2015, 2015; Haegens et al., 2011; Jensen et al., 2014; Lundqvist et al., 2024; Mathewson et al., 2011; Osipova et al., 2008; Samaha et al., 2017; Sauseng et al., 2009; Staresina et al., 2016; Voytek et al., 2010). For example, it is easier to elicit the motor evoked potential (MEP) with transcranial magnetic stimulation (TMS) when alpha power at the motor cortex is low (Sauseng et al., 2009). Moreover, alpha/beta power correlates with lower thresholds for TMS-induced phosphenes (Romei et al., 2008; Samaha et al., 2017). Therefore, spindles overlapping with task-related alpha/beta reductions (Petzka et al., 2022) may reflect a mechanism whereby the expression of spindles is flexible and tracks ‘hot spots’ of cortical excitement during prior wakefulness.

If spindles track regional cortical excitability, it should not matter whether cortical excitement is induced endogenously through particular learning tasks (Clemens et al., 2005, 2006; Petzka et al., 2022) or exogenously through experimental brain stimulation. Here we test this notion by directly exciting local brain regions with transcranial direct current stimulation (tDCS) and tracking the expression of spindles as well as SOs in subsequent sleep. Anodal-tDCS has been shown to increase cortical excitability in local circuits for over 90 minutes after application (Agboada et al., 2019; Ho et al., 2016; Stagg et al., 2018; Stagg & Nitsche, 2011; Vignaud et al., 2018; Woods et al., 2016), allowing us to boost excitation before sleep and observe its effects on regional sleep spindles. Moreover, anodal-tDCS is thought to facilitate synaptic plasticity by increasing calcium uptake by N-methyl-D-aspartate (NMDA) receptors and reducing inhibitory gamma-aminobutyric acid (GABA) levels (Bachtiar et al., 2015; Nitsche et al., 2003; Stagg et al., 2018). Interestingly, pharmacologically influencing NMDA receptors and GABA modulates spindles (Fernandez & Lüthi, 2020; Jacobsen et al., 2001; Mednick et al., 2013). Therefore, tDCS may induce physiological aftereffects at cortical sites which in turn modulate subsequent spindle expression. Finally, to examine the potential effects of modulating local spindles on memory consolidation, we used a unilateral visuomotor finger-tapping task thought to be regionally localised to right motor cortex (Ambrus et al., 2016; Buch et al., 2021; Iwane et al., 2023; Witt et al., 2008) to observe potential changes in motor learning behaviour from before to after sleep (Boutin et al., 2018; Fogel et al., 2017).

Our results indicate that cortical excitability generated by tDCS can modulate the rate of spindles at targeted sites. Importantly, we did not find an effect of excitatory tDCS on the distribution of SOs. These results suggest spindle topographies are neither hard-wired nor random but are influenced by pre-sleep cortical excitability and can be modified with external stimulation.

## Methods

### Participants

We collected data from 21 volunteers. One participant was excluded because of a technical issue and another due to inability to sleep, resulting in a final sample size of 19 (9 female; age, M = 25.00 years, SD = 4.37). All participants were right-handed according to the Edinburgh Handedness Inventory (Oldfield, 1971) and were not proficient in playing a keyboard instrument. Participants were also screened to ensure they did not work nightshifts in the last month, were not pregnant, were non-smokers, did not take sleep-altering medication, and did not have a history of neurological or psychiatric disorders. Participants were asked to get up an hour earlier than usual on the day of the session. They were also asked to refrain from consuming alcohol or caffeine on the night before or the day of the sessions, respectively. All participants self-reported an ability to nap in the daytime. Written informed consent was provided by all participants at the beginning of each session and participants were compensated a total of 150 GBP for their time. Protocols were approved by the local ethics committee (Central University Research Ethics Committee #R83711/RE002).

### Procedure

Participants engaged in two day-long experimental sessions from 9:00–17:00, at least 5 days apart. **Figure 1A** illustrates the timeline of an experimental session. After providing written consent and being screened for MRI and tDCS safety, the participant washed and dried their hair and changed into scrubs. The experimenter then applied the EEG cap. At 10:00 the participant started with the computerized ‘localiser’ and visuomotor finger-tapping tasks which lasted approximately 60 minutes (see **Tasks** for details).

**Figure 1.**
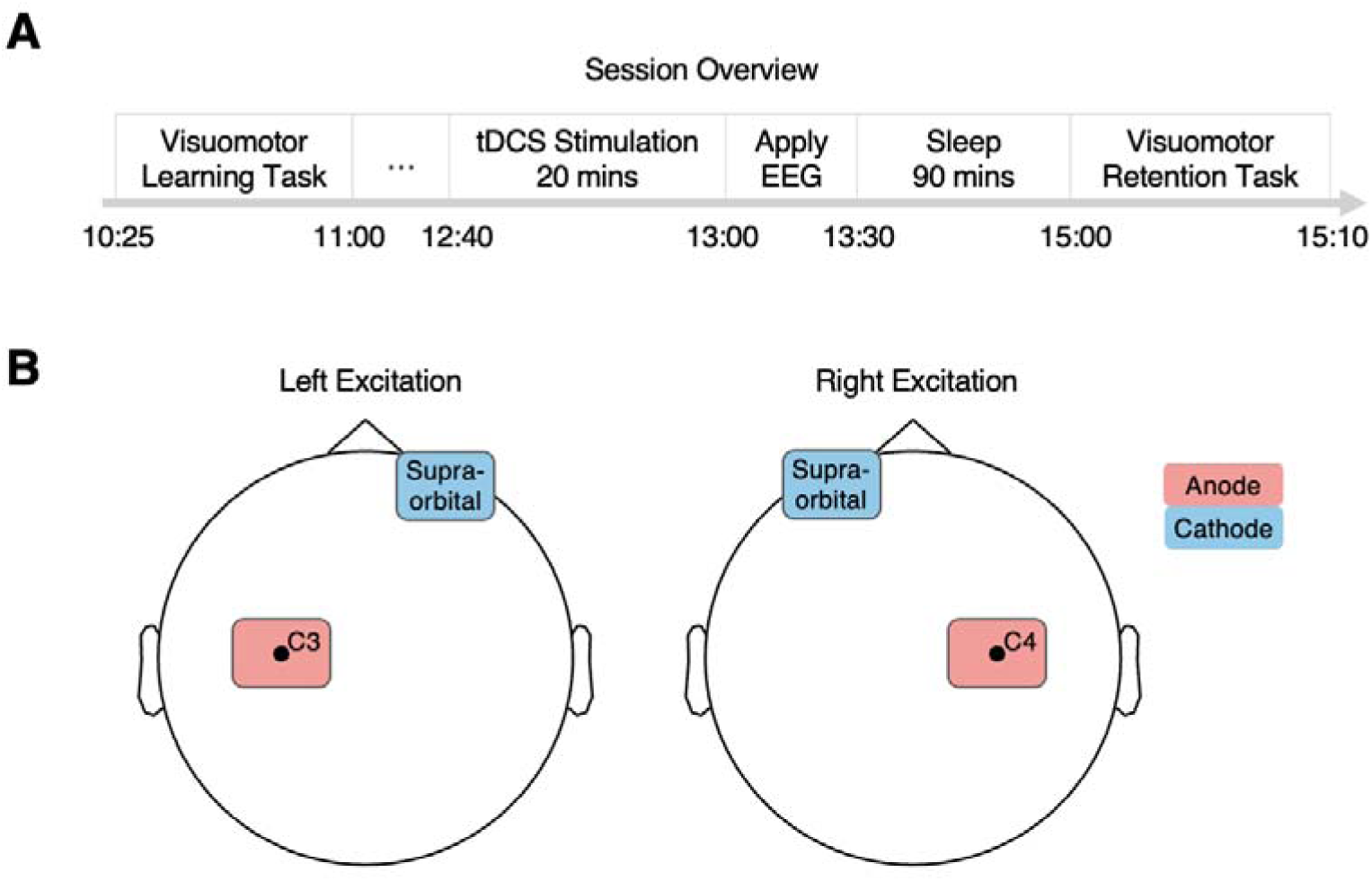
Session overview and tDCS montages. **A)** Overview of a session’s schedule. Each participant conducted two sessions at least 5 days apart; one session with left excitation and one with right excitation (order counterbalanced across participants). **B)** tDCS montages for the left and right excitation conditions. Stimulation was set to 1mA for 20 minutes. A simulation of the distribution of voltage induced by the tDCS montages is shown in **Supplementary Figure 2**.

After completing the computerized tasks, the experimenter removed the EEG cap, and the participant washed and dried their hair. At 11:30 AM, the participant underwent structural and functional MRI scans lasting 45 minutes. From 12:15 to 12:30, participant and experimenter ate lunch whilst watching a nature documentary. The documentary was randomly selected for each session and each participant. At 12:30, the experimenter applied the tDCS pads which took 10 minutes. A total of 1.5 hours elapsed between completing the tasks and the application of tDCS, allowing residual task-related activation to subside before stimulation.

Next, the tDCS was switched on and ran at 1mA for 20 minutes, during which time the experimenter and participant continued passively watching the nature documentary. After the stimulation, the tDCS pads were removed, the scalp was cleaned with alcohol, and the experimenter reapplied the EEG cap. Once the EEG cap was applied, the participant got into bed and wore earplugs and an eye mask to sleep. The experimenter left the room, giving the participant 90 minutes to sleep.

At 15:00, the experimenter woke the participant and gave them five minutes to overcome sleep inertia. The participant then completed three more blocks of the visuomotor finger-tapping task to assess motor skill post-sleep. After completing the retest of the finger-tapping task, the EEG cap was removed, the participant washed and dried their hair before having another set of MRI scans which lasted approximately one hour. At 17:00 the participant changed back into their clothes and was thanked for their time.

### Tasks

The tasks were run using MATLAB 2021 (Mathworks, 2012) and Psychtoolbox (Brainard, 1997). The participant used an iMac keyboard and a 23.8-inch colour screen (1920×1080 resolution) which was positioned approximately 30 cm away. To keep the participant from looking at their fingers during the tasks, a cardboard box was placed over their left hand.

The first task was a visual localiser in which images were shown on the screen. The participant was instructed to press the down-arrow key if the same image was presented twice in a row. The images in each session were unique and consisted of two faces, two objects, two scenes, and two words. Each image was presented 55 times in total and was repeated twice in a row five times. Presentations lasted 1 sec, with a 0.5 sec fixation period and a random intertrial-interval jitter of between −0.1 and 0.1 secs. There were self-paced pauses after 75 image presentations which participant skipped by pressing the space bar.

Next, the participant learned the unilateral visuomotor finger-tapping task. First, they received instructions regarding the task design. The task consisted of blocks which contained a tapping phase and a resting phase (see **Supplementary Figure 1**). During the tapping phase, four of the images from the localiser task would be presented in a sequence. Each image belonged to one category (face, object, scene, or word) and each category was linked to a number key at the top of the keyboard (1-face, 2-object, 3-scence, 4-word). The participant used their left hand to press the number keys, with one key assigned to each finger. The goal was to press the corresponding key as quickly as possible when an image was presented. For example, an image of George Clooney would require the participant to press key “1” since it is linked to the category “face”. After pressing “1”, the next image would be presented, which is a desert scene prompting the participant to press key “3” (linked to scene). This would continue for 30 secs, pressing keys in the sequence as quickly and accurately as possible.

The participant was explicitly instructed that there was a four-part sequence which would repeat for the entire session. The sequences were “1-3-2-4” and “4-2-3-1” which are mirrors of each other, to match complexity. The image stimuli and tapping sequence used in session 1 and session 2 were counterbalanced between participants. After the tapping phase, the participant received feedback, which was displayed for 3 secs, on how many full four-part sequences (“1-3-2-4” or “4-2-3-1”) they had pressed. They were instructed to maximise the number of full correct sequences over the course of the blocks.

After the tapping phase and feedback, there was a resting phase in which the participant performed a distraction task where they needed to count the number of times a fixation cross changed shades of grey. This resting phase kept the participant focused whilst allowing their left hand to rest. After the resting phase, the participant entered the number of fixation cross changes using their right hand and a separate keyboard from the one used in the tapping phase. That way their left hand remained in the same position, on the number keys for the entire task. The next block began once the number of fixation changes was entered.

The visuomotor finger-tapping task before the nap consisted of 23 blocks. Each block was made up of tapping phase, feedback, and resting phase. The participant was instructed to try as hard as they could throughout the learning blocks to perform well in the final test blocks which were identical to the learning blocks. The resting phase of the 10^th^ and 20^th^ blocks were extended to five minutes to allow the participant’s left hands to significantly rest. Additionally, before starting with the learning blocks, the participant performed a practice block with different images and a different sequence. In total, the visuomotor finger-tapping task lasted approximately 40 minutes.

### tDCS Application

The coordinates of the stimulation sites (C3 and C4) were located with a measuring tape during the initial EEG capping. Before stimulation, the sites were marked using a red marker to guide the tDCS application. The tDCS system was a NeuroConn DC-Stimulator Plus with 5 cm x 7 cm pads. After parting hair and cleaning the scalp with rubbing alcohol, we used Ten20 conductive paste to adhere the pads to the scalp as well as rubber bands. **Figure 1B** depicts how the anodal pad was placed at C3 or C4 with a vertical orientation while the cathodal pad was placed on the contralateral supraorbital cortex with a horizontal orientation. The tDCS ran at 1mA for 20 minutes whilst the participant and experimenter continued watching a nature documentary. We chose this montage because it has been shown to influence synaptic and cortical excitability in ipsilateral motor cortex (Agboada et al., 2019; Bachtiar et al., 2015; Vignaud et al., 2018; Woods et al., 2016). Since there was no sham condition, participants were informed they would receive stimulation in both sessions and that they may experience a tingling sensation around the electrode pads. It was not possible to blind the experimenter to whether C3 or C4 was being stimulated because the conditions required different pad-placements. However, participants were blind to the importance of the pad-placements in each condition.

### EEG Data Recording

EEG recordings were conducted using a 64-contact Brain Products ActiChamp system (*Brain Products UK*) recording at 1000 Hz. The cap used a standard 10-20 layout which was customised to have two electrooculography (EOG), two electromyography (EMG), and two mastoid contacts for sleep scoring. AFz was used as the ground electrode and Fz was the online reference. The impedance of the contacts was brought below 20 kOhm with Abralyt HiC gel. Lic2 electrode cream was used to attach the two EMG electrodes to the skin under the chin. Finally, bandage gauze was applied around the EEG cap to secure it during sleep.

### Sleep Scoring

Sleep scoring was performed on the raw data, split into in 30 sec epochs, by two deep learning models, Somnobot (*SomnoBot*, n.d.) and YASA (Krieter et al., 2020). A researcher, blind to the conditions, visually inspected the epochs and resolved discrepancies between the classifications of the two models according to the American Academy of Sleep Medicine guidelines (Berry, R. B. et al., 2020).

### EEG Preprocessing and Event Detection

EEG data were analysed in MATLAB (Mathworks, 2012) using Fieldtrip functions (Oostenveld et al., 2011). The recordings were down-sampled to 200 Hz and the mastoid, EOG, and EMG contacts were removed. Artefacts were detected on each contact individually as events with an absolute amplitude over 2.5 SD from the mean or an absolute amplitude over 100 mV. Artefacts were padded with 1 sec on either side and contacts whose signal was more than 10% artefactual were interpolated from neighbouring contacts using “ft_channelrepair” with the “weighted interpolation” method. After interpolating bad contacts, all contacts were re-referenced to the mean signal of all contacts.

Custom scripts based on established detection algorithms (Ngo et al., 2020) were used to detect spindle and SO events. Only non-artefactual data during N2 and N3 epochs were considered for event detection. To detect spindles, the EEG signal was first bandpass filtered between 12-16 Hz and the envelope was smoothed with a 0.2 sec root mean square kernel. A spindle event was identified when the smoothed envelope had an amplitude between 1.5-5 SD from the mean of the eligible data and had a duration between 0.5-3 secs. For the more conservative threshold criteria used in **Supplementary Figure 6**, the detection threshold was 1.75-5 SD from the mean.

To detect SOs, the signal was filtered to retain frequencies between 0.3 and 1.25 Hz. All points where the signal crossed zero were identified, and the candidate SO duration was measured as the interval between the two consecutive positive-gradient zero crossings. Candidate events lasting between 0.8 and 2 secs were then selected, and their amplitude was evaluated (both trough and trough-to-peak amplitude). Events were classified as SOs if both trough and trough-to-peak amplitudes were above the mean plus 1.5 SD of all candidate events.

### Behavioural and EEG Statistical Analysis

We used MATLAB (Mathworks, 2012) and the Fieldtrip toolbox (Oostenveld et al., 2011) to analyse the data. Data and analysis scripts to reproduce the main results are available online at https://osf.io/8cnxf/.

In the visuomotor finger-tapping task, RTs were excluded which were below 0.05 secs and above 0.5 secs, to remove accidental presses and outliers. The first block of one session was excluded from the RT analyses because the participant did not press a correct key with an RT under 0.5 secs. Spindle and SO rates were calculated per session as the number of events divided by the total non-artefactual time spent in N2 or N3 (given in events per minute).

Cluster-based permutation tests were performed using “ft_timelockstatistics” in the Fieldtrip toolbox, using 1000 random permutations, a cluster-alpha threshold of *p* < .05 and a significance threshold of *p* < .05 (two-tailed). The data distribution was presumed to be normal, though this assumption was not verified.

To explore the effect of power spectra on skill retention (**Supplementary Figure 7**), we first calculated the fast Fourier transform of the signal for frequencies 0-30 Hz (steps of 0.5 Hz) in 4 sec windows (50% overlap) of N2 and N3 sleep. Next, we binned the power into 2 Hz bins and ran linear-mixed effect models (LMEs) with the equation ‘Retention ∼ Session + Stimulation Site + Spectral Power + (1|Participant Number)’, where retention was the average performance in the three blocks after sleep minus the three blocks before sleep. A separate LME was run for each frequency bin and contact combination (15 bins x 57 contacts = 855 combinations), whilst controlling for anodal-tDCS stimulation site and session number.

## Results

### Polysomnography and spindle events

Participants slept for an average of 78.74 mins (SD = 12.13) in each session. This included an average of 60.68 mins of N2 or N3 sleep (N2, M = 41.62 mins, SD = 12.95; N3, M = 19.07 mins, SD = 12.32). **Figure 2A** depicts a hypnogram from an example session with time spent in different sleep stages. LME models showed neither session number (1 vs. 2) nor stimulation condition (right vs. left excitation) had a significant effect on the total time or proportion of time spent in N2 and N3. Further summary statistics of sleep stages are detailed in **Supplementary Table 1**.

**Figure 2.**
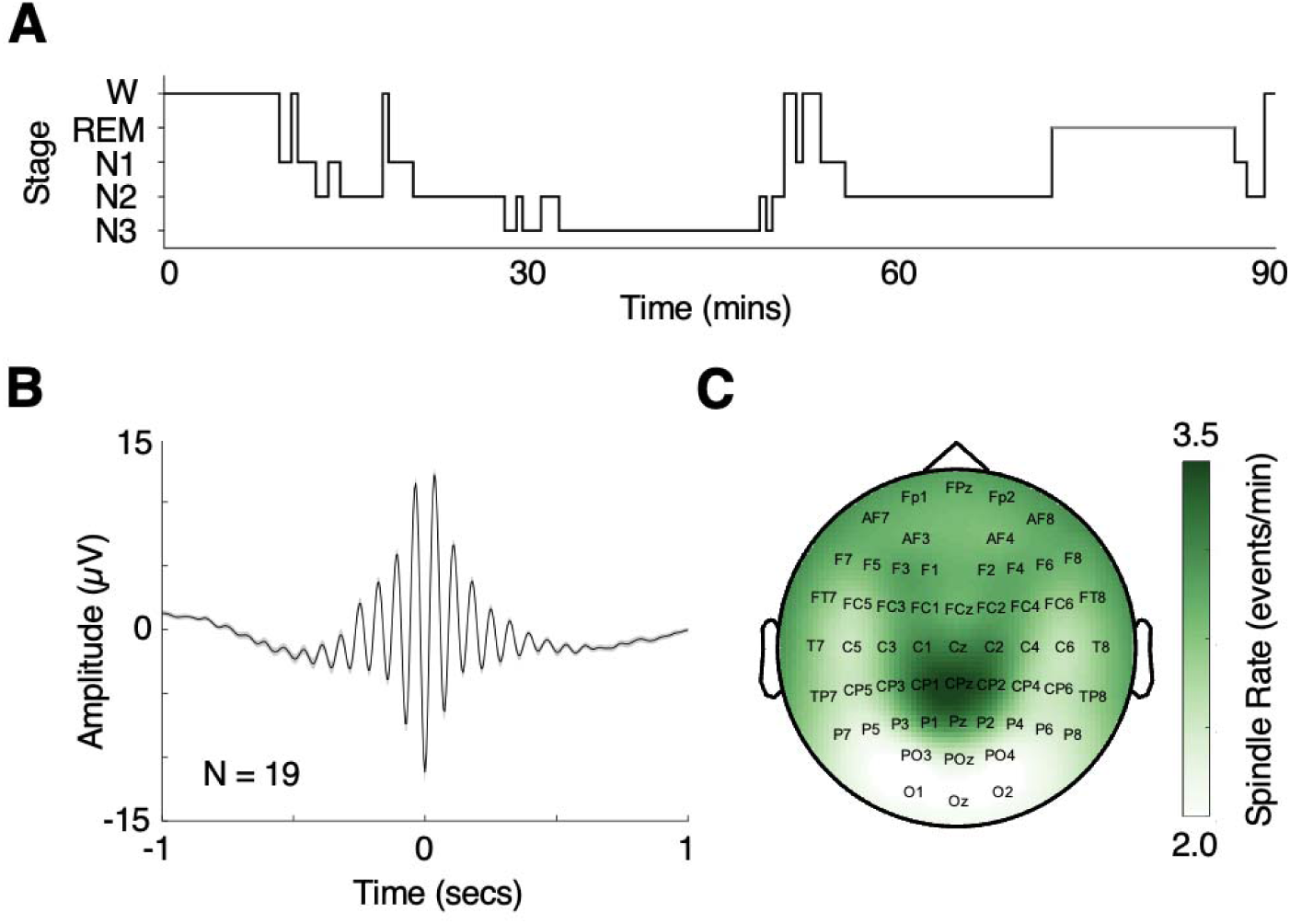
Sleep stages and spindle events. **A)** Example hypnogram showing time spent in different sleep stages. **B)** Grand average spindle from Cz contacts. Spindles were aligned to their largest trough and baseline corrected using the mean of the 2 sec window, before averaging across sessions (N = 2) and across participants (N = 19). Shading represents ±1 SEM. **C)** Average spindle rate topography. The topographies were averaged across left and right excitation sessions before averaging across participants.

Figure 2B shows the waxing and waning nature of the grand average spindle detected at Cz across participants. The spindle rate topography averaged across both excitation conditions was most dense around Cz and parietal areas (Figure 2C) and the spindle rates averaged 2.15 and 2.16 events/min at the C3/C4 target sites respectively (**Supplementary Table 2**).

### Targeted tDCS stimulation increased local spindle rates

To quantify the effect of targeted, pre-sleep excitatory stimulation on subsequent spindle rates, we conducted a paired-samples t-test to compare the C4 minus C3 spindle rate contrast in sessions with left and right excitatory stimulation. If spindle rates are greater at excited sites, we would expect the C4 (right) minus C3 (left) contrast to be greater when C4 (right) is excited, compared to when C3 (left) is excited. Figure 3A confirms this, showing a significant difference in C4 minus C3 spindle rate contrasts between right excitation (M = 0.02, SD = 0.27) and left excitation conditions (M = −0.17, SD = 0.37), *t*(18) = −2.55, *p* = .020, *d* = −0.58, two-tailed). The same t-test was not significant for SO rate contrasts (**Supplementary Figure 3A**).

**Figure 3.**
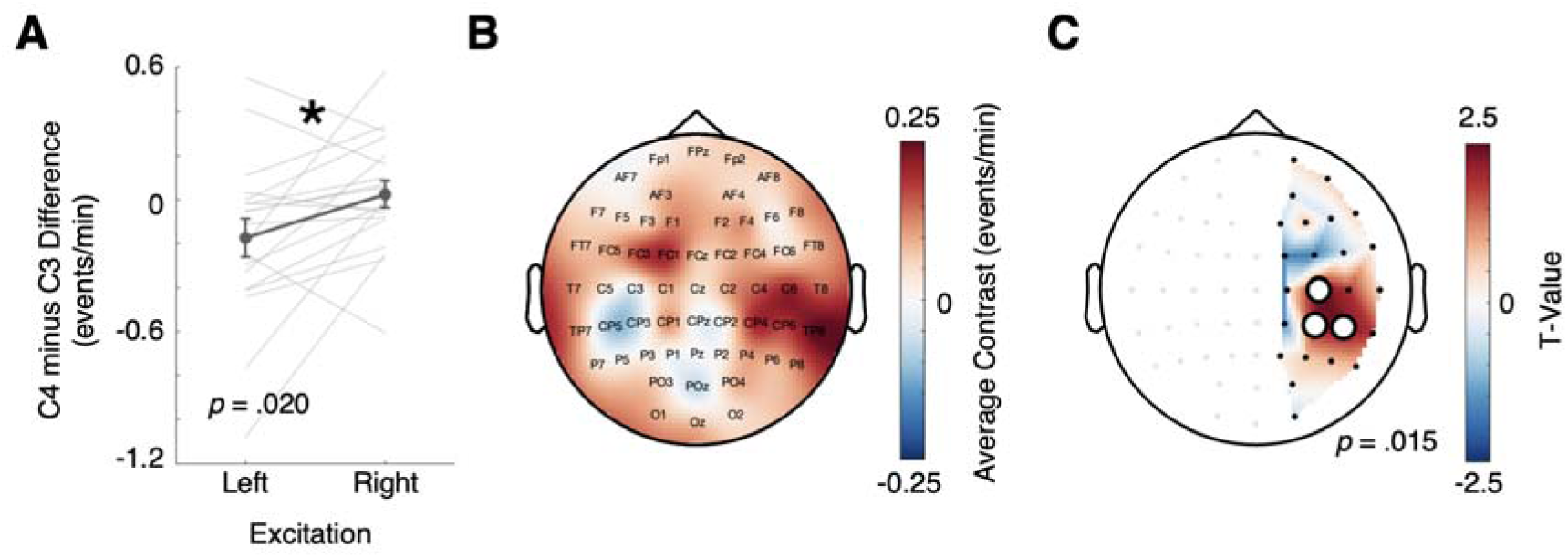
Spindle rates were influenced by lateralised tDCS stimulation. **A)** Significant contrast between spindle rates at C3 and C4 target sites in left and right excitation conditions. Lines represent individual participants. Error bars represent ±1 SEM. **B)** Average right excitation spindle rate topography minus left excitation spindle rate topography contrast across participants (N=19). **C)**. Right excitation minus left excitation contrast values from left hemisphere contacts were subtracted from right hemisphere contacts to create a measure of excitation-dependent spindle rate lateralisation. White circles depict a significant cluster, using cluster-based permutation tests.

To test the effect of all aforementioned factors on event rate, we used a 3-way repeated-measures ANOVA (factor 1 = event type (spindle/SO), factor 2 = stimulation condition (left/right excitation), factor 3 = EEG contact (C3/C4)). Results showed a significant main effect of event type (*F*(1,18) = 12.13, *p* = .003, η*_p_^2^* = .40), with spindles having a greater rate than SOs (spindles, M = 2.73, SD = 0.36; SOs, M = 2.44, SD = 0.27; **Supplementary Figure 4**). Importantly, there was a significant 3-way interaction between event type, stimulation condition, and EEG contact (*F*(1,18) = 5.59, *p* = .030, η*_p_^2^* = .24), confirming the effect of stimulation condition on event rate at the target contacts is specific to spindles and not SOs. Together, these results show spindle rates at targeted sites were significantly modulated using excitatory pre-sleep tDCS.

Next, we further investigated the regional specificity of the effect of excitatory stimulation on spindle rates. **Figure 3B** shows the topography of the contrast in spindle rates between sessions with right excitatory stimulation (anode on C4) and left excitatory stimulation (anode on C3). To test the regional specificity of the effect of targeted lateral stimulation, we subtracted the right excitation minus left stimulation contrast of left hemisphere contacts from their right hemisphere mirror pairs (e.g., CP3 from CP4) and compared this lateralised contrast value to zero. **Figure 3C** shows a significant positive cluster including C3/C4, CP3/CP4, and CP5/CP6 (cluster-based permutation test, *p* = .015). The same analysis and test methods did not show any significant effects of lateralised tDCS on regional SO rates (**Supplementary Figures 3B-C**), nor did it affect the amplitudes or durations of the spindle events. These results demonstrate the boosting effect of targeted pre-sleep tDCS stimulation on spindle rate is localised to the target region and does not carry over to SOs.

To test the robustness of the findings, we repeated the analysis with z-scored spindle rates. Before taking the contrasts, we normalised spindle rates across the scalp EEG contacts within each session. **Supplementary Figure 5A** shows the paired-samples t-test with significant differences in z-scored C4 minus C3 spindle rate contrasts between left excitation (M = −0.35, SD = 0.67) right excitation (M = −0.01, SD = 0.65), *t*(18) = −2.18, *p* = .042, *d* = −0.50, two-tailed. **Supplementary Figures 5B-C** illustrate the corresponding right minus left excitation condition contrast, showing a significant cluster at C3/C4 and CP5/CP6 (*p =* .043). As a final control, we increased the threshold for detecting spindle events during event detection from mean plus 1.5 SD to mean plus 1.75 SD, thus making the detection criteria more conservative (**Supplementary Figure 6A**). Repeating the analysis with only the spindle events fulfilling the stricter criteria again led to similar results. The paired-samples t-test illustrated in **Supplementary Figure 6B** shows a significant difference in the C4-C3 contrast between left excitation (M = −0.15, SD = 0.27) and right excitation (M = 0.00, SD = 0.26) conditions, *t*(18) = − 2.72, *p* = .014, *d* = −0.62, two-tailed. **Supplementary Figure 6C** displays the hemisphere contrasts of the right minus left excitation contrasts with a significant positive cluster at C3/C4, CP5/CP6, and TP7/TP8. Together, these results suggest that the effect of targeted tDCS on spindle rate lateralisation is robust to normalisation and to more conservative thresholds for spindle events.

### Lateralised excitatory stimulation did not significantly influence skill retention after sleep in a unilateral finger-tapping task

Sleep spindles have been linked to memory consolidation during sleep (Bergmann et al., 2012; Boutin et al., 2018; Fogel et al., 2017; Petzka et al., 2022; Schreiner et al., 2021). By comparing the performance in the unilateral visuomotor finger-tapping task, we were able to examine if excitatory tDCS stimulation to the left or right motor cortex influenced skill retention after sleep and how this might be mediated by changes in local sleep spindles. To this end, we used two LME models to examine the effect of block and session on average reaction time (RT; secs) of correct key presses (model 1) and the number of correct four-part sequence responses (model 2), with participant included as a random intercept. Results indicated a significant main effect of block in both models, suggesting that RT decreased across blocks (*β* = −0.004, SE = 0.000, *t*(870) = −23.10, *p* < .001) and the number of correct four-part sequence responses increased across blocks (*β* = 0.332, SE = 0.017, *t*(871) = 19.00, *p* < .001). This reduction in RT and increase in correct sequence responses with successive blocks provides evidence of learning over time, as participants became faster and more accurate with practice. There was also a main effect of being in the second session on RT (*β* = −0.011, SE = 0.002, *t*(870) = −5.91, *p* < .001), indicating differences in RT between sessions. Similarly, the number of full sequences was greater in the second session (*β* = 0.761, SE = 0.232, *t*(871) = 3.65, *p* < .001), suggesting performance was generally better in the second session.

To evaluate whether tDCS stimulation condition influenced motor skill retention in the visuomotor finger-tapping task, we tested for an interaction between performance metrics (mean RT of correct key presses and mean number of full correct sequence responses) across the three penultimate trials of the learning task and the mean performance metrics across the three trials following sleep, similar to Fogel and colleagues (Fogel et al., 2017). This analysis was conducted using two 2 × 2 repeated-measures ANOVAs. Performance was significantly better in the trials pre-sleep, compared to trials post-sleep (RT: *F*(1,18) = 11.30, *p* = .003, η*_p_^2^* = .61; correct sequences: *F*(1,18) = 15.01, *p* = .001, η*_p_^2^* = .41). However, there was no main effect of stimulation condition (RT: *F*(1,18) = 0.05, *p* = .834, η*_p_^2^*= .00; correct sequences: *F*(1,18) = 1.86, *p* = .189, η*_p_^2^* = .05) and no significant interaction effect between pre-sleep and post-sleep performance and right-versus left-excitatory tDCS stimulation conditions (RT: *F*(1,18) = 0.18, *p* = .677, η*_p_^2^*= .01; correct sequences: *F*(1,18) = 0.79, *p* = .387, η *^2^*= .04). Therefore, the results do not provide evidence that tDCS stimulation condition influences skill retention and memory consolidation across sleep in the unilateral visuomotor finger-tapping task.

Although there was no effect of stimulation site on skill retention after sleep, it is possible that other oscillatory signatures were linked to skill retention of the visuomotor task. To explore this, we extracted spectral power at each contact from 0-30 Hz, in 2 Hz bins. We used an LME model to test the effect of spectral power on skill retention for each combination of frequency bin and contact, whilst controlling for stimulation site and session number (see **Methods** for details). **Supplementary Figure 7** shows an exploratory trend of 6-10 Hz power affecting skill retention across primarily frontal and fronto-central contacts. Positive t-values and negative t-values indicate more retention when considering the number of correct sequences or the average RT per block, respectively. Note that these results are exploratory and the tests for significance were not corrected for multiple comparisons. This suggests, if anything, 6-10 Hz oscillations, and not localised spindles, may contribute to skill retention in the visuomotor task employed here and provides insight as to why modulating spindles at C3/C4 did not affect skill retention after sleep in our paradigm.

## Discussion

We set out to test whether the local expression of sleep spindles is sensitive to cortical excitability during wakefulness, exogenously induced via tDCS. Our results demonstrate excitatory tDCS can modulate regional spindle rates in humans. Specifically, anodal-tDCS applied before a nap increased the number of spindles at the target site compared to the contralateral homologous region. The effect was specific to the target region, significantly influencing spindle rates at the stimulation sites (C3/C4) and neighbouring centroparietal sites (Figure 3A-C). SO rates have also been shown to track learning-related sites (Huber et al., 2004; Nir et al., 2011; Vyazovskiy et al., 2011), however, the effect of anodal-tDCS was specific to spindles in our current study. The significant interaction shows tDCS was greater for spindles rates than SO rates (**Supplementary Figure 4**), ruling out that stimulation had a global/broadband effect. These results indicate spindle topographies are flexible and are affected by pre-sleep cortical excitability, which can be directly influenced using non-invasive brain stimulation.

The spatial specificity in our results aligns with previous work showing that spindle expression tracks the topography of cortical excitability, induced endogenously via pre-sleep learning tasks. While previous work (Petzka et al., 2022) showed spindle topographies adapt to match task-induced activation patterns, our results extend this by revealing spindles also track regional excitability induced through experimental exogenous stimulation. Together these findings corroborate the notion that the deployment of sleep spindles is adaptive and biased towards regions that are excited before sleep.

Previous studies that tried to modulate spindles with slow-oscillating or alternating transcranial stimulation yielded mixed results (Antonenko et al., 2013; Barham et al., 2016; Cellini & Mednick, 2019; Del Felice et al., 2015; Eggert et al., 2013; Koo et al., 2018; Ladenbauer et al., 2016, 2017; Lustenberger et al., 2016; Marshall et al., 2004, 2006, 2011; Mednick et al., 2013; Park et al., 2023; Sahlem et al., 2015). These studies attempted to experimentally stimulate spindles during sleep by inducing pulsing or alternating currents. Rather than treating the brain as a resonator during sleep, we here used tDCS before sleep to modify the excitability of the underlying cortical circuits with constant excitatory stimulation using a well-tested protocol (Agboada et al., 2019; Stagg et al., 2018; Woods et al., 2016). Specifically, the tDCS protocol used here has been shown to modulate cortical excitability and increase the amplitude of motor evoked potentials in motor cortex up to 90 minutes after stimulation (Agboada et al., 2019). The success of this method in modulating the topography of spindles suggests targeting the underlying synaptic and physiological states of cortical regions (Bachtiar et al., 2015; Chittajallu et al., 1998; Stagg et al., 2018) and thus “tagging” them for later endogenous spindle deployment may be more fruitful than attempting to entrain specific bursts with their frequency profiles.

Despite successfully modulating local spindles around motor areas, we did not find evidence for behavioural influences of excitatory tDCS on a left-handed finger-tapping task which we assumed to locally recruit right motor areas around C4. Therefore, we cannot draw conclusions on the effect of modulating spindles on regionally specific motor memory consolidation during sleep. Firstly, it is possible that the effect of local spindles on regional memory consolidation is not linear, and influencing the expression of spindles may have a more complex effect on learning retention behaviour. Second, our task may not have been optimally designed to benefit from local increases in spindle rates. **Supplementary Figure 7** shows an exploratory trend of fronto-central 6-10 Hz oscillations predicting visuomotor skill retention after sleep, suggesting that if at all, oscillatory signals outside the fast spindle range may contribute to skill consolidation during sleep. Finally, an ultimate measure of motor-learning after sleep is difficult to obtain, since the test blocks themselves constitute more practice and participants thus continue to learn during testing. This issue of continual learning limits how many ‘trials’ can enter the post-sleep test, making results vulnerable to outliers. Alternative paradigms like standard declarative memory tasks would allow for more trials during testing, albeit perhaps at the cost of clearly circumscribed cortical hotspots as elicited through finger-tapping.

Nevertheless, our findings have several potential applications. The ability to guide spindles using targeted non-invasive brain stimulation may be valuable for targeted neurorehabilitation, particularly when modulation of regional plasticity is needed. Brain-computer interfaces during sleep which look to modify activity patterns in the sleeping brain may be able to use targeted excitation to bias local sleep oscillations and thus affect sleep quality and/or memory performance. Additionally, this work provides a foundation for investigating the effect of targeted stimulation before sleep on the expression and characteristics of sleep oscillations. Future work may explore other sleep oscillations, stimulation protocols, and application methods to refine our understanding of the malleability of spatiotemporal dynamics of sleep oscillations.

In conclusion, our results demonstrate that pre-sleep cortical excitability can shape the topographical expression of sleep spindles in humans, suggesting a flexible system which is sensitive to exogenous as well as endogenous excitation of cortical circuits. These findings lay the groundwork for future work to investigate regionally specific, sleep-dependent memory processes and to develop targeted interventions for manipulating regional sleep oscillations in research and the clinic.

## Acknowledgements

We thank Mathew Kollamkulam for his extensive support whilst piloting and calibrating the study.

## Supplementary Material

**Supplementary Figure 1.**
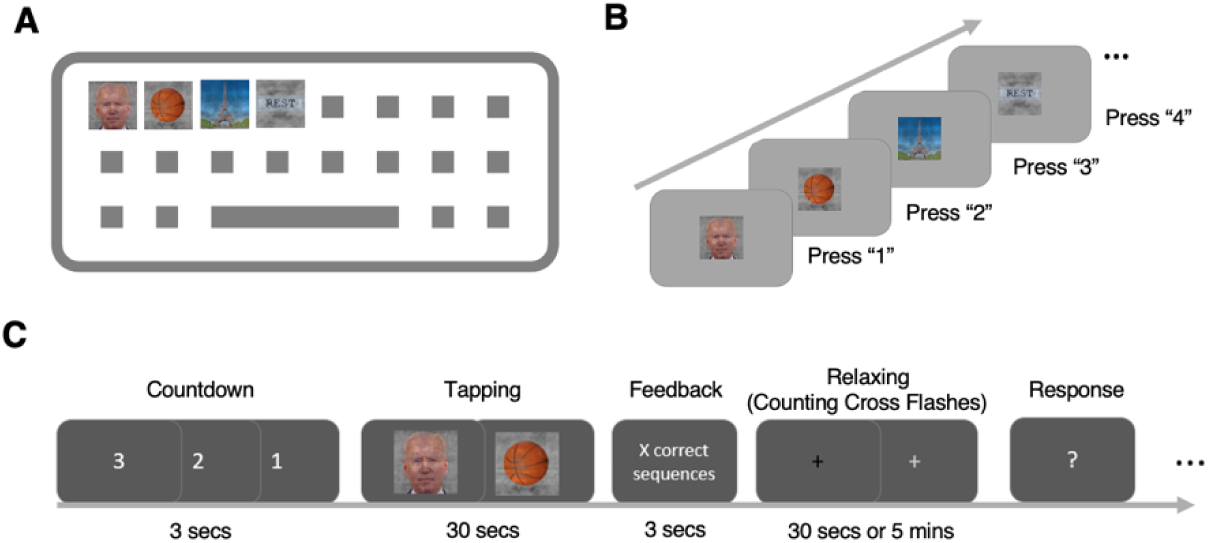
Overview of visuomotor finger-tapping task. **A)** An image category was assigned to each of the four number keys in the top row of the keyboard. The participant received explicit instructions to use one finger per key. The assignment was always 1-Face, 2-Object, 3-Scene, 4-Word, although the images were unique to each session and were counterbalanced across subjects. **B)** Pressing the image category key would progress the task to the next image. **C)** Overview of a block. After a countdown, the participant tapped for 30 secs before receiving feedback on the number of correct four-part sequences they achieved. They then relaxed their hand and counted the number of times the fixation cross changed brightness for 30 secs (5 mins after block 10 and block 20) and finally responded how many flashes they counted with their right hand on a separate keyboard. There were 23 blocks in total during the learning task and three blocks after sleep.

**Supplementary Figure 2.**
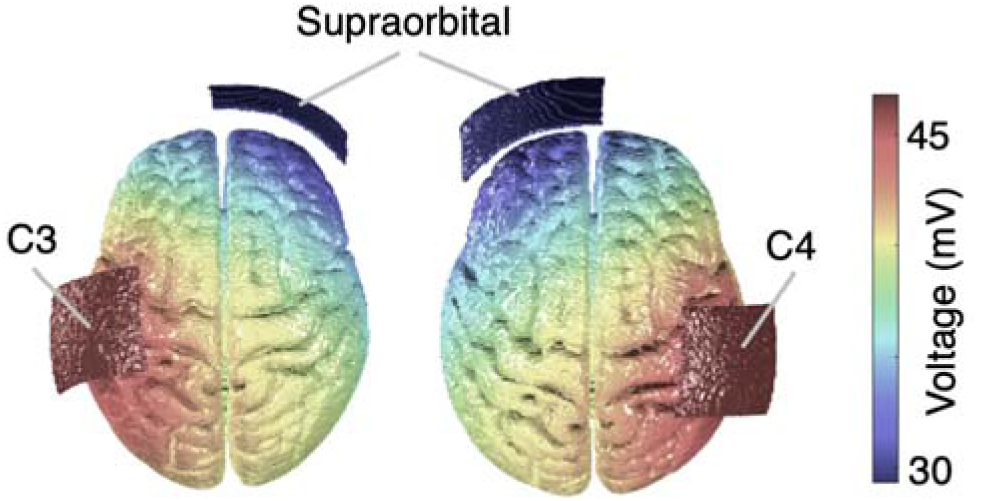
Simulation of tDCS voltage effects. **A)** Simulation using ROAST (Huang et al., 2019) of the distribution of voltage caused by 1mA tDCS stimulation for 20 mins during left excitatory and right excitatory stimulation. The simulation was conducted using a template brain.

**Supplementary Figure 3.**
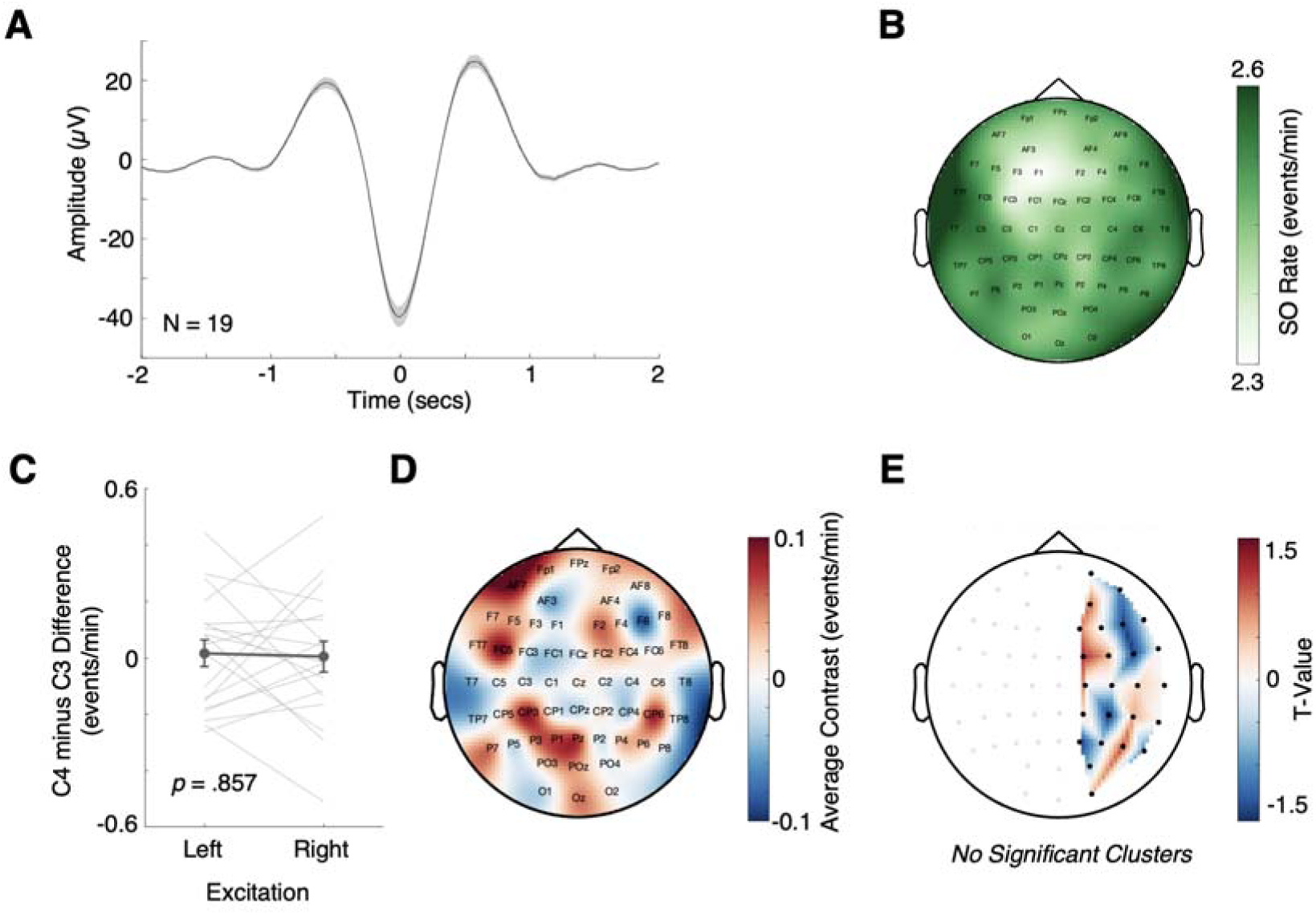
No evidence of tDCS stimulation influencing the lateralisation of SO rates. **A-E)** Same as **Figure 2B-C** and **Figure 3A-C** with SO rates. Lines represent individual participants. Error bars represent ±1 SEM. T-test was paired-sample and two-sided. Cluster-based permutation tests were tested as p<.05 significance.

**Supplementary Figure 4.**
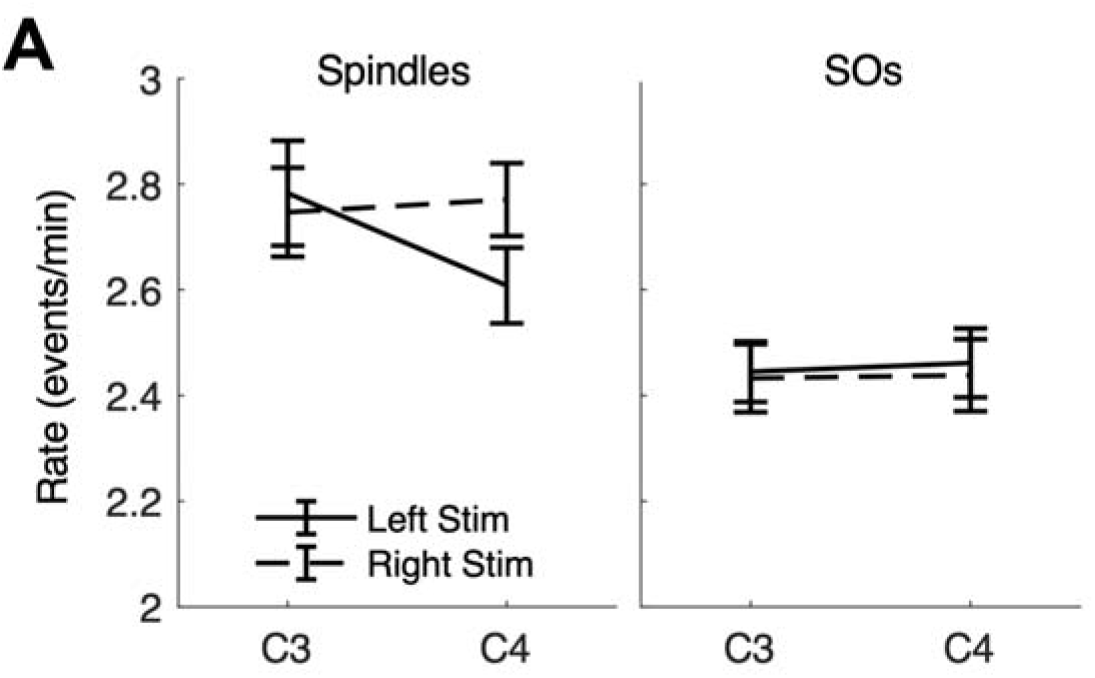
3-way ANOVA and interaction between event type, stimulation condition, and contact. **A)** The mean event rates across subjects, split by event type, contact, and stimulation condition. Error bars indicate ±1 SEM. The main effect of event type was significant, F(1,18) = 12.13, p = .003, ηp^2^ = .40, with spindles having a greater rate than SOs (spindles, M = 2.73, SD = 0.36; SOs, M = 2.44, SD = 0.27). The 3-way interaction between event type, stimulation condition, and contact was significant (F(1,18) = 5.59, p = .030, ηp^2^ = .24).

**Supplementary Figure 5.**
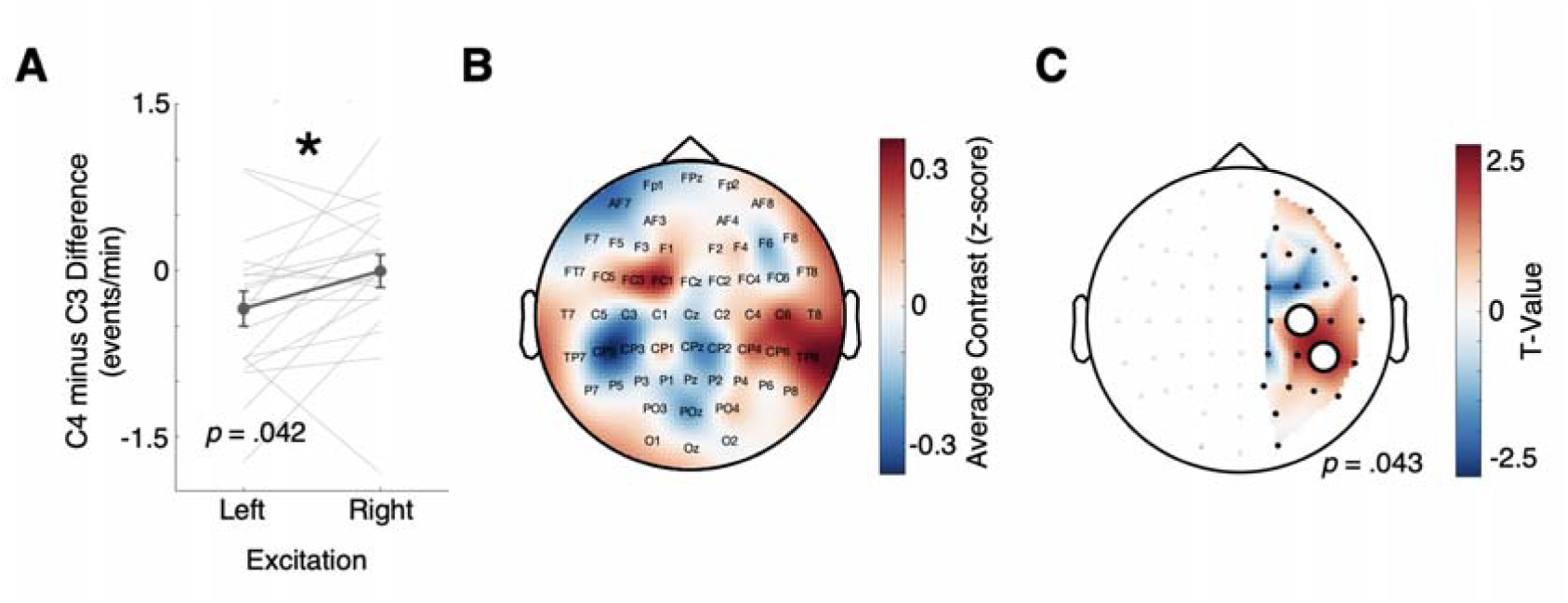
Z-scored spindle rates were influenced by lateralised tDCS stimulation. **A-C)** Same as **Figure 3** with z-scored spindle rates (normalised within each session before subtracting and creating contrasts). Error bars represent ±1 SEM. Lines represent individual participants. White circles highlight significant clusters, p < .05.

**Supplementary Figure 6.**
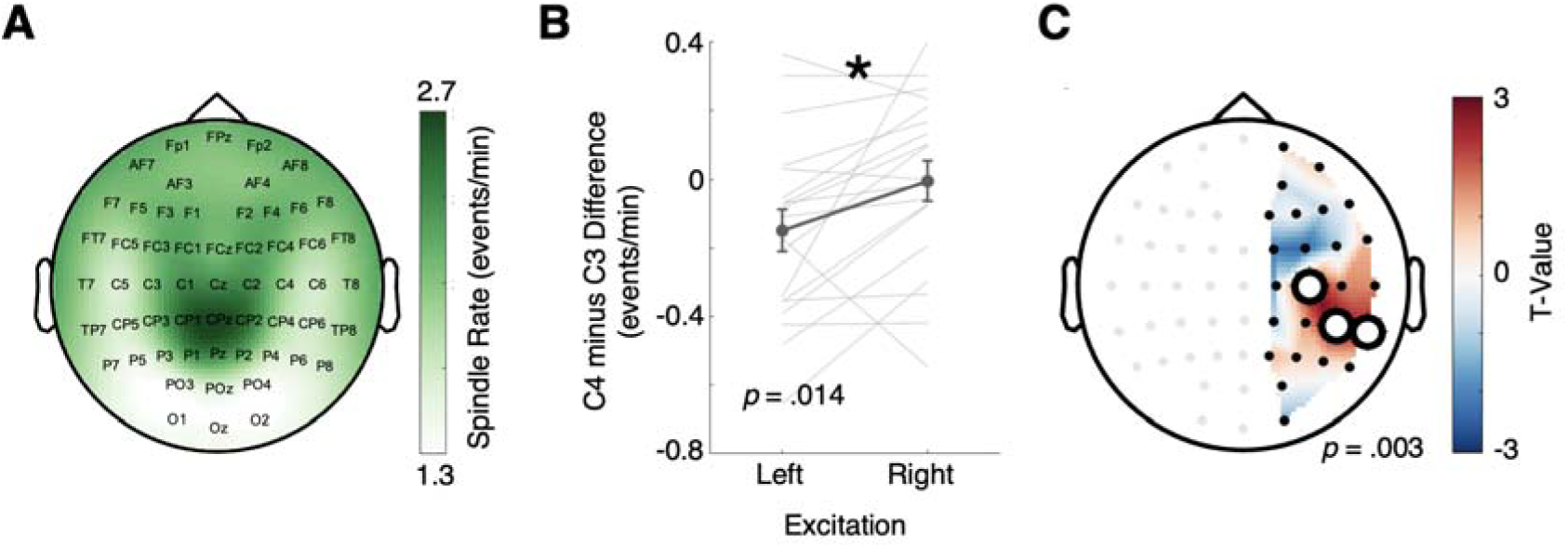
Spindle rates with more conservative thresholds were influenced by lateralised tDCS stimulation. **A-C)** Same as **Figures 2C, 3A, & 3C** with spindle rates using a detection threshold of mean + 1.75 SD (rather than mean + 1.5 SD). Lines represent individual participants. Error bars represent ±1 SEM. White circles highlight significant clusters, p < .05.

**Supplementary Figure 7.**
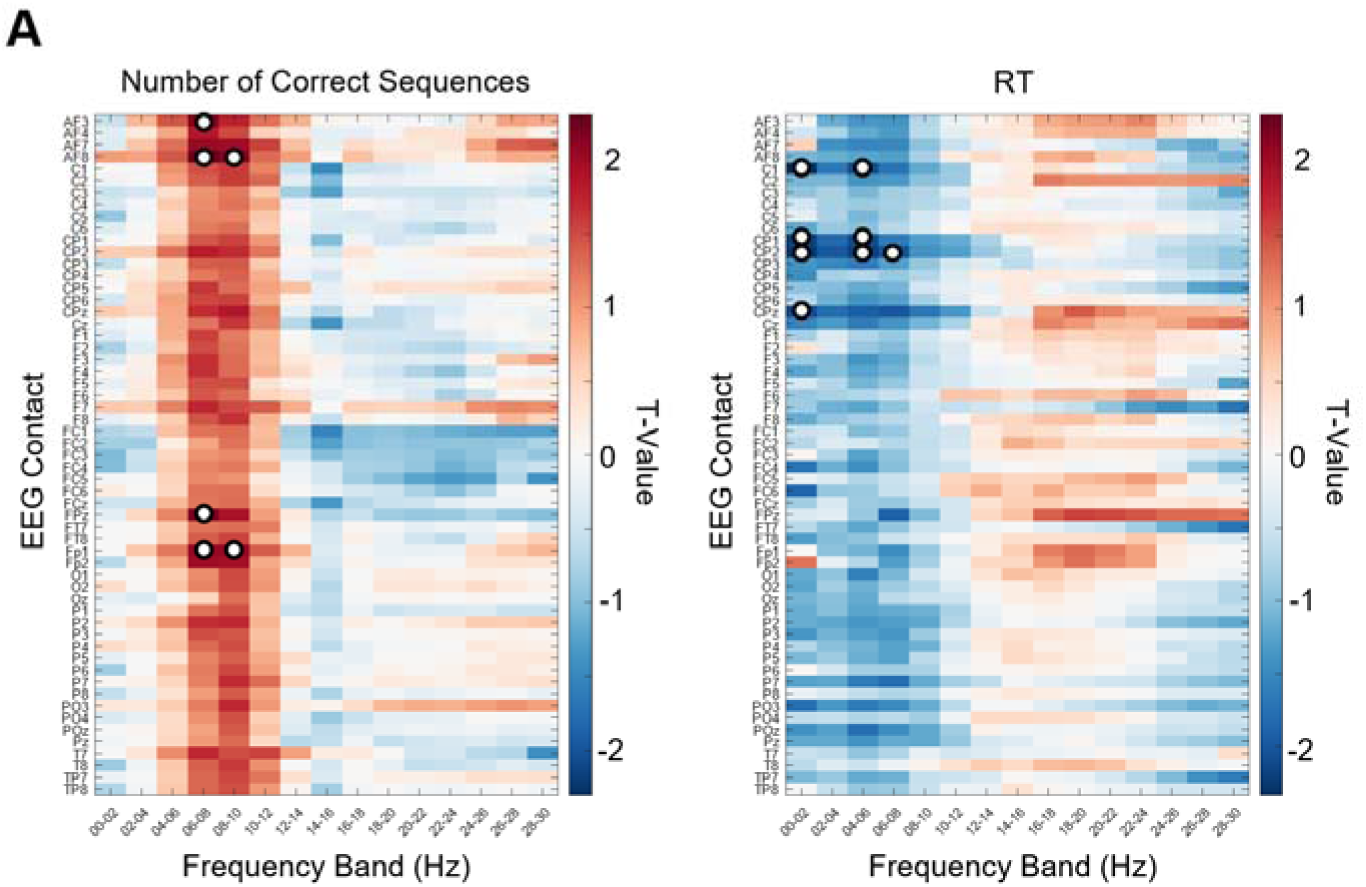
Exploration of relationship between spectral power in N2 / N3 sleep and visuomotor skill retention after sleep. **A)** The t-values for the effect of spectral power on skill retention from exploratory LME models with the equation ‘Retention ∼ Session + Stimulation Site + Spectral Power + (1|Participant Number)’. One LME model was employed for each EEG contact-frequency bin combination (see **Methods** for details). Positive and negative t-values indicate better retention after sleep for the number of correct sequences and RT, respectively. White circles represent significance (p < .05), not corrected for multiple comparisons.

**Supplementary Table 1.** Summary sleep statistics.

| Metric | Mean | SD | Min | Max |
| --- | --- | --- | --- | --- |
| Time N1 (mins) | 12.41 | 8.70 | 2.50 | 44.00 |
| Time N2 (mins) | 41.62 | 12.95 | 18.00 | 63.50 |
| Time N3 (mins) | 19.07 | 12.32 | 0.00 | 45.50 |
| Time N2/3 (mins) | 60.68 | 16.69 | 23.00 | 91.50 |
| Time REM (mins) | 5.64 | 8.16 | 0.00 | 29.00 |
| Total Sleep Time (mins) | 78.74 | 12.13 | 49.50 | 100.00 |
| Proportion N1 | 0.17 | 0.13 | 0.03 | 0.66 |
| Proportion N2 | 0.53 | 0.14 | 0.26 | 0.78 |
| Proportion N3 | 0.24 | 0.15 | 0.00 | 0.55 |
| Proportion N2/3 | 0.76 | 0.15 | 0.34 | 0.97 |
| Proportion REM | 0.07 | 0.10 | 0.00 | 0.35 |

**Supplementary Table 2.** Summary spindle statistics for C3, C4, and Cz.

| Metric | Contact | Mean | SD | Min | Max |
| --- | --- | --- | --- | --- | --- |
| Number of Spindles | C3 | 165.82 | 45.81 | 67 | 268 |
|  | C4 | 162.00 | 44.54 | 62 | 258 |
|  | Cz | 191.61 | 61.35 | 58 | 350 |
| Spindle Rate<br>(events/min) | C3 | 2.76 | 0.40 | 2.15 | 3.92 |
|  | C4 | 2.69 | 0.31 | 2.16 | 3.40 |
|  | Cz | 3.16 | 0.53 | 1.96 | 4.24 |

